# No balance between glutamate+glutamine and GABA+ in visual and motor cortices of the human brain

**DOI:** 10.1101/2021.03.02.433628

**Authors:** Reuben Rideaux

## Abstract

Theoretical work, supported by electrophysiological evidence, asserts that a balance between excitation and inhibition (E/I) is critical for healthy brain function. In magnetic resonance spectroscopy (MRS) studies, the ratio of excitatory (glutamate) and inhibitory (γ-aminobutyric acid, GABA) neurotransmitters is often used as a proxy for this E/I balance. Recent MRS work found a positive correlation between GABA+ and Glx (glutamate+glutamine) in medial parietal cortex, providing validation for this proxy and supporting the link between the E/I balance observed in electrophysiology and that detected with MRS. Here we assess the same relationship, between GABA+ and Glx, in primary visual and motor cortices of resting male and female human participants. We find moderate to strong evidence that there is no positive correlation between these neurotransmitters in either location. We show this holds true when controlling for a range of other factors (i.e., demographics, signal quality, tissue composition, other neurochemicals) and regardless of the state of neural activity (i.e., resting/active). These results show that there is no brain-wide balance between excitatory and inhibitory neurotransmitters and indicates a dissociation between the E/I balance observed in electrophysiological work and the ratio of MRS-detected neurotransmitters.

## INTRODUCTION

Balanced excitatory and inhibitory activity (i.e., E/I balance) is a canonical feature in models of healthy brain (Marín, 2012; Shadlen & Newsome, 1994; Van Vreeswijk & Sompolinsky, 1996; Vogels & Abbott, 2009), which is supported by a large body of empirical evidence, primarily from electrophysiological experiments. Diverse external stimulation, such as a bar of light (Anderson, Carandini, & Ferster, 2000), a whisker deflection (Wilent & Contreras, 2005), a brief tone (Wehr & Zador, 2003), or an odour (Poo & Isaacson, 2009), all evoke concurrent excitatory and inhibitory activity, which is thought to bestow neurons with sharper tuning properties (Higley & Contreras, 2006; Wilent & Contreras, 2005). The phenomenon can even be observed during periods of spontaneous activity and is likely responsible for maintaining equilibrium throughout the brain (Okun & Lampl, 2008). By contrast, E/I imbalance is implicated in central pathologies including epilepsy (Bradford, 1995; Olsen & Avoli, 1997), autism spectrum disorder (Chao et al., 2010; Markram & Markram, 2010; Rubenstein & Merzenich, 2003; Vattikuti & Chow, 2010), and schizophrenia (Kehrer, Maziashvili, Dugladze, & Gloveli, 2008).

Magnetic resonance spectroscopy (MRS) can be used to measure *in vivo* concentrations of primary excitatory (glutamate) and inhibitory (γ-aminobutyric acid, GABA) neurotransmitters within the brain. It is common practice in MRS studies to report the ratio of these neurotransmitters as an index of E/I balance. This proxy has been used to study a range of healthy brain functions including decision-making (Bezalel, Paz, & Tal, 2019), visual learning (Shibata et al., 2017), volitional control (Koizumi, Lau, Shimada, & Kondo, 2018), memory (Bang et al., 2018; Takei et al., 2016), and visual contrast sensitivity (Ip, Emir, Parker, & Bridge, 2018). It has also been used to investigate other neuroimaging signals, such as the default mode network activity (Gu, Hu, Chen, He, & Yang, 2019; Kapogiannis, Reiter, Willette, & Mattson, 2013). In the context of theories relating neuropsychiatric conditions to E/I imbalance, GABA and glutamate or Glx (a complex comprising glutamate and glutamine) concentrations have been identified as a potential biomarker for autism spectrum disorder and schizophrenia (Brown, Singel, Hepburn, & Rojas, 2013; Egerton et al., 2012; Horder et al., 2013; Smesny et al., 2015).

A recent MRS study tested the validity of using the ratio of excitatory and inhibitory neurotransmitters as a proxy for E/I balance (Steel, Mikkelsen, Edden, & Robertson, 2020). The researchers found that the concentration of Glx and GABA+ (GABA + co-edited macromolecules) in medial parietal cortex was positively correlated across a large cohort of healthy participants. This evidence suggests an association between electrophysiological and MR spectroscopic observations of E/I balance. However, if these measurements reflected a common neural phenomenon (i.e., E/I balance), we would expect to observe a positive correlation between GABA+ and Glx at other cortical regions where concomitant excitatory and inhibitory activity has been reported. Thus, to assess the regional specificity of this association, we tested for a relationship between GABA+ and Glx in two different cortical locations: early visual and motor cortices. These brain regions are an ideal testbed of this association, as numerous electrophysiological studies have demonstrated their E/I balance (Anderson et al., 2000; Dehghani et al., 2016; Maffei & Turrigiano, 2008; Priebe & Ferster, 2005; Wilent & Contreras, 2005; Xue, Atallah, & Scanziani, 2014).

In contrast to Steel et al. (2020), we find no evidence for a relationship between GABA+ and Glx across participants in either visual or motor cortices. Post-hoc Bayesian analyses reveal moderate to strong evidence in favour of the null hypothesis. We further show that the same (null) result is found in visual cortex even during visual stimulation, indicating that the discrepancy between the results from medial parietal cortex and visual/motor cortices cannot be explained by differences in overall neural activity. These findings show that there is no canonical brain-wide balance between GABA+ and Glx, and suggest that the balance reported in medial parietal cortex may be the exception, rather than the rule. Moreover, the results indicate a dissociation between the E/I balance observed using other techniques (e.g., electrophysiology) and the ratio of excitatory and inhibitory neurotransmitters detected with MRS.

## METHODS

### Data collection

Legacy data from previous studies were combined. In these studies, male and female human participants underwent MR spectroscopic acquisition targeting visual (n=58; 31 women; mean age=24.4; age range=19-40; Rideaux, 2020; Rideaux, Goncalves, & Welchman, 2019; Rideaux & Welchman, 2018), motor (n=50; 24 women; mean age=24.3; age range=19-36; Rideaux & Welchman, 2018) cortices while at rest. During the acquisition, the lights in the room were turned off and participants were instructed to close their eyes. In addition to this, data from visual cortex of participants (n=31; 17 women; mean age=24.9; age range=19-40; Rideaux et al., 2019) who received visual stimulation during acquisition was analysed as a control. All data were collected on the same vendor at the same site. All participants were screened for contra-indications to MRI prior to data collection. All experiments were conducted in accordance with the ethical guidelines of the Declaration of Helsinki and were approved by the University of Cambridge ethics committee and all participants provided informed consent.

### Data acquisition

Magnetic resonance scanning was conducted on a 3T Siemens Prisma equipped with a 32-channel head coil. Anatomical T1-weighted images were acquired for voxel placement with an MP-RAGE sequence. For detection of GABA+, spectra were acquired using a MEGA-PRESS sequence (Mescher, Merkle, Kirsch, Garwood, & Gruetter, 1998): TE=68 ms, TR=3000 ms; 256 or 400 transients of 2048 data points were acquired, 16 water-unsuppressed transients were additionally acquired; a 14.28 ms Gaussian editing pulse was applied at 1.9 (ON) and 7.5 (OFF) ppm. Water suppression was achieved using variable power with optimized relaxation delays (VAPOR; Tkáč and Gruetter, 2005) and outer volume suppression. Automated shimming with 3D GRE, followed by manual shimming, was conducted to achieve approximately 12 Hz water linewidth. All MRS data were analysed from their DICOM format.

Spectra were acquired from locations targeting visual and motor cortices (**Fig. 1a**). The voxel targeting visual cortex (n=40, 3×3×2 cm; n=29, 2.5×2.5×2.5 cm) was positioned medially in the occipital lobe with the lower face aligned with the cerebellar tentorium. The voxel targeting motor cortex (3×3×2 cm) was centred on the ‘hand knob’ area of the precentral gyrus and aligned to the upper surface of the brain in the sagittal and coronal planes. The coordinates of the voxel location were used to draw a mask on the anatomical T1-weighted image to calculate the fractional volume of grey matter, white matter, and cerebrospinal fluid within each voxel. Tissue segmentation was performed using the Statistical Parametric Mapping toolbox for MATLAB (SPM12, www.fil.ion.ucl.ac.uk/spm; Ashburner & Friston, 2005).

**Figure 1.**
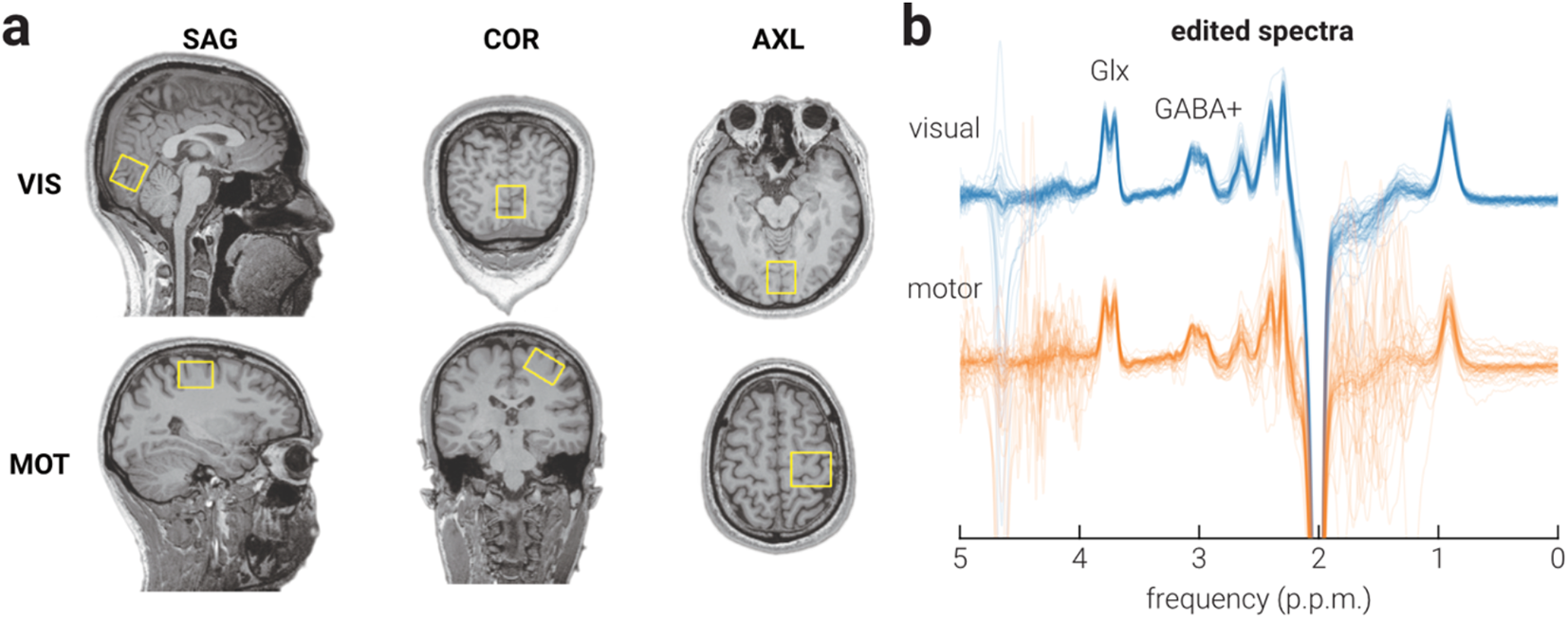
Voxel locations and average edited spectra. **a**) Sagittal (SAG), coronal (COR), and axial (AXL) views of representative MRS voxel placement for visual (VIS) and motor (MOT) cortices on a T1-weighted structural image. **b**) Average edited spectra for resting visual and motor cortices.

The conditions of the ‘visual stimulation’ control data acquisition were the same as those described in the resting visual cortex dataset, with the exception that participants viewed black and white random-dot kinetograms during the acquisition. In particular, participants performed an attentionally demanding vernier task at fixation while an annulus comprising black and white dots, with a fan-shaped three-dimensional profile, was presented around fixation. Participants were required to monitor fixation for the lines used to perform the vernier task, which appeared at pseudorandom intervals throughout the scan. The random-dot kinetograms were presented for 1.8 s and separated by 0.2 s inter-stimulus-intervals throughout the scan. For a detailed description of the stimulus and presentation procedure, see the ‘mixed-polarity’ condition in Rideaux et al. (2019).

### Data processing

Spectral pre-processing and quantification was conducted in MATLAB using a combination of Gannet v3.1 (Edden et al., 2014) and in-house scripts. Prior to alignment, subspectra were zero-filled to a spectral resolution of 0.061 Hz/point and 3-Hz exponential line-broadening was applied. Frequency and phase parameter estimates were obtained by modelling the total creatine (tCr; creatine and phosphocreatine) signal, then these parameters were used to align subspectra to a common frequency and phase. Note that we found that same pattern of results when Spectral Registration was used to align spectra. Estimates of signal area and full-width at half-maximum (FWHM) were also obtained, and subspectra (and their corresponding ON/OFF subspectra) with parameter estimates >3 standard deviations (s.d.; the default cut-off applied in Gannet) from the mean within a scan were omitted from further analysis.

Total creatine and total N-acetyl aspartate (tNAA; N-acetyl aspartate and N-acetyl aspartyl glutamate) signal intensity were determined by fitting a Lorentzian model to the average OFF spectra at 3.0 ppm and 2.0 ppm, respectively. The average ON and OFF spectra were subtracted to produce the edited spectrum (**Fig. 1b**), from which GABA+ (3 ppm) and Glx (3.8 ppm) signal intensity were modelled. Glx and GABA+ were fit using a double-Gaussian model. Water signal intensity was determined by fitting a Lorentzian model to the average water-unsuppressed spectra at 4.9 ppm. All neurochemical signal intensities were calculated as the area of the fitted peak(s) and expressed in institutional units (i.u.) using the unsuppressed water signal as an internal concentration reference. The assumed relaxation and density parameters of water and GABA are described in (Mikkelsen et al., 2019). The assumed longitudinal relaxation times of Glx, tCr, tNAA, and Cho were 1.23 s, 1.35 s, 1.41 s, and 1.19 s, respectively (Posse et al., 2007). The assumed transverse relaxation times of these metabolites were 0.18 s, 0.15 s, 0.25 s, 0.21 s, respectively (Ganji et al., 2012).

### Statistical analyses

Statistical analyses were conducted in MATLAB (The MathWorks, Inc., Matick, MA). Prior to correlational analyses, we screened the data by omitting (n=3, all from motor cortex group) participants with GABA+ or Glx values >2.5 s.d. from the mean, in line with (Steel et al., 2020). We sought to determine whether the concentration of GABA+ and Glx were related across different brain regions. To test this, we computed the Pearson correlation between GABA+ and Glx for all spectra. Next, to test whether the relationship GABA+ and Glx could be explained by confounding factors we systematically regressed out the influence of factors that could potentially account for the observed relationship using a linear mixed effects model. We then computed the Pearson correlation between the residuals after controlling for the confounding factors. We reasoned that if the residual values remained correlated, the relationship between GABA+ and Glx held when controlling for these confounding variables. To establish the influence of confounding factors, we considered each factor separately and calculated the significance of the change in correlation by comparing z-scored correlation coefficients. Power analyses were conducted in GPower (www.gpower.hhu.de; Erdfelder, Faul, & Buchner, 1996). Bayesian analyses were conducted in JASP (www.jasp-stats.org; Wagenmakers et al., 2018).

## RESULTS

### No balance between GABA+ and Glx

We found no evidence for a relationship between GABA+ and Glx across participants in either visual (*r*_58_=-.09, *P*=.493) or motor cortices (*r*_47_=-.05, *P*=.762). Power analyses, based on the correlation between GABA+ and Glx in medial parietal cortex found by Steel et al. (2020) (r=.40), showed that the sample sizes tested here had a 94% and 89% probability of detecting the same relationship if it existed in visual and motor cortices, respectively. There are multiple factors that may influence the quantification of GABA+ and/or Glx with MRS. It is possible that a correlation between these neurotransmitters in visual and/or motor cortices was masked by an additional moderating factor. To test this possibility, we systematically assessed the relationship between GABA+ and Glx after regressing out a range of possible moderating factors. The results of these analyses are shown in **Table 1**.

**Table 1.** List of confounding factors tested.

| Confounding factor | Correlation of GABA+ and Glx residuals |  |
| --- | --- | --- |
|  | visual | motor |
| age and gender | $r=-.09, p=.494$ | $r=-.01, p=.925$ |
| frequency drift | $r=-.08, p=.563$ | $r=-.02, p=.879$ |
| water linewidth | $r=-.09, p=.504$ | $r=-.04, p=.804$ |
| tNAA linewidth | $r=-.13, p=.334$ | $r=-.002, p=.991$ |
| GABA+ fit error | $r=-.10, p=.466$ | $r=-.05, p=.763$ |
| Glx fit error | $r=-.11, p=.417$ | $r=-.05, p=.727$ |
| grey matter | $r=-.10, p=.467$ | $r=-.03, p=.830$ |
| tNAA concentration | $r=-.30, p=.021$ | $r=-.32, p=.029$ |
| tCr concentration | $r=-.22, p=.096$ | $r=-.30, p=.040$ |
| Cho concentration | $r=-.29, p=.026$ | $r=-.30, p=.040$ |
| water concentration | $r=-.11, p=.442$ | $r=-.16, p=.269$ |

### Age and gender

Previous MRS work indicates that demographic characteristics such as age and gender are related to the concentration of GABA+ and Glx. That is, in adults, the concentration of GABA+ and Glx tends to be higher in males (O’Gorman, Michels, Edden, Murdoch, & Martin, 2011), and decreases with age (Cassady et al., 2019; Gao et al., 2013; Sailasuta, Ernst, & Chang, 2008). However, we found no significant difference in the relationship between GABA+ and Glx in either visual or motor cortices as a result of controlling for age and gender (**Fig. 2b**; visual: *z*_57_=0.001, *p*=.999; motor: *z*_47_=0.15, *p*=.884).

**Figure 2.**
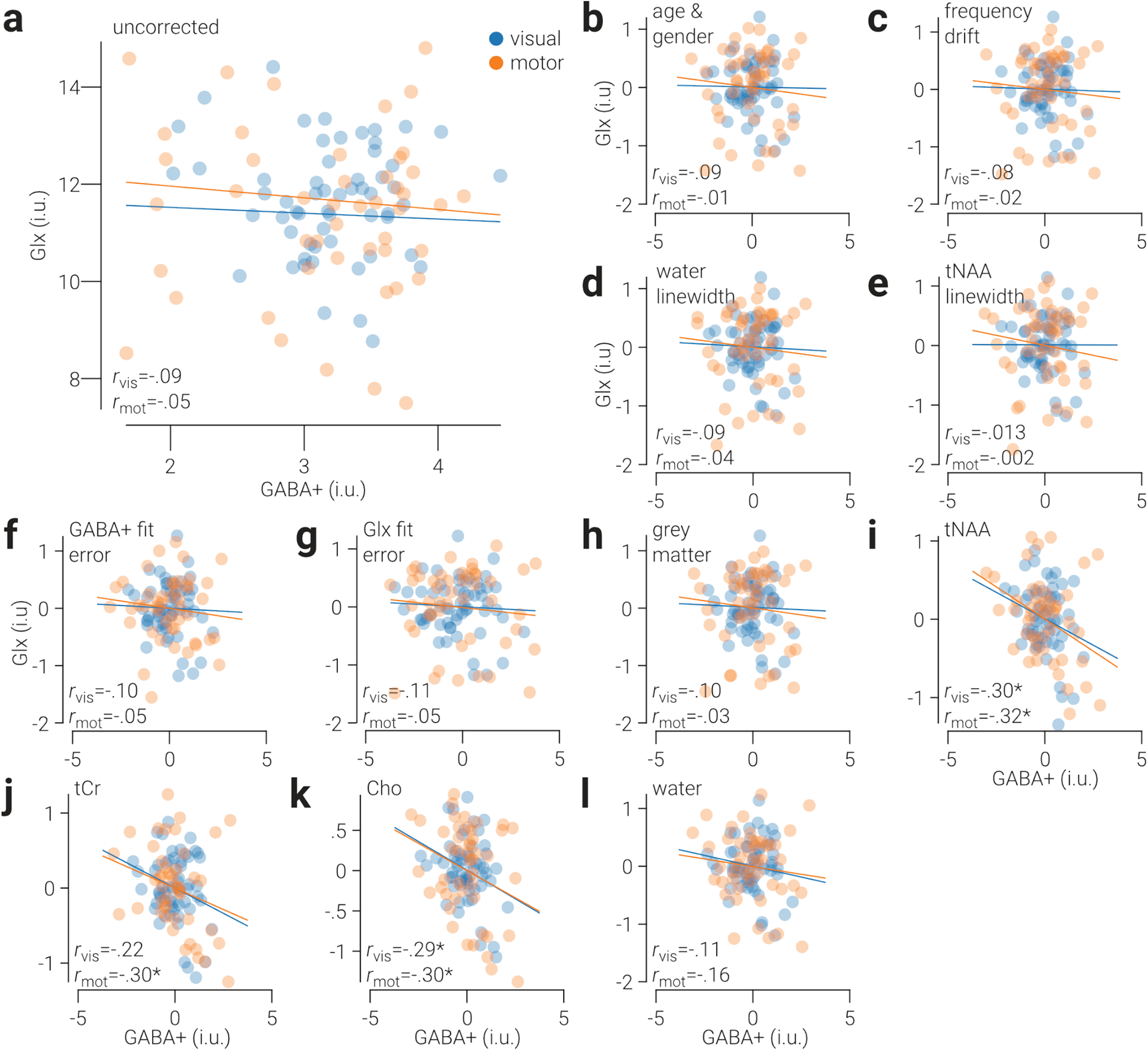
No balance between GABA+ and Glx in visual and motor cortices. **a**) Glx concentration as a function of GABA+ concentration in early visual and motor cortices. **b-g**) The same as (**a**), after controlling for **b**) age and gender, **c**) frequency drift, **d**) water linewidth, **e**) tNAA linewidth, **f**) GABA+ fit error, **g**) Glx fit error, **h**) grey matter, **i**) tNAA concentration, **j**) tCr concentration, **k**) Cho concentration, and **l**) water concentration. Glx and GABA+ values are referenced to water and expressed in institutional units (i.u.). Lines indicate best linear fit, and cyan and orange colours indicate data from visual and motor cortices, respectively. Asterisks indicate correlation coefficients where *p*<.05.

### Spectral quality

As both GABA+ and Glx signals were quantified from the same spectra, differences in signal quality may have obscured their relationship. That is, low signal quality could result in systematic misestimation of both GABA+ and Glx, producing a spurious correlation that masks a putative non-spurious relationship between these neurochemicals.

To assess this possibility, we regressed out the differences in three measures of signal quality (frequency drift, tNAA linewidth, and water linewidth) before re-testing the relationship between GABA+ and Glx. Frequency drift can occur from participant head motion or field gradient heating/cooling and can impact quantification of GABA+ and Glx by altering the efficiency with which their signals are edited (Harris et al., 2014). The editing efficiency of GABA+ decreases (modestly) with frequency drift in either direction, whereas that of Glx can increase or decrease depending on the drift direction (Harris et al., 2014). Thus, frequency drift could produce either a positive or negative spurious correlation between GABA+ and Glx. However, we found no significant difference in the relationship between GABA+ and Glx in either visual or motor cortices as a result of controlling for frequency drift (**Fig. 2c**; visual: *z*_57_=0.08, *p*=.940; motor: *z*_47_=0.11, *p*=.916). The linewidth of metabolite peaks provides an indication of signal quality, as larger linewidths are associated with poorer spectral acquisition, e.g., due to participant head motion or inefficient shimming (Harris et al., 2014). However, we found no significant change in the correlation between GABA+ and Glx as a result of controlling for either water (**Fig. 2d**; visual: *z*_57_=0.01, *p*=.990; motor: *z*_47_=0.04, *p*=.969) or tNAA linewidth (**Fig. 2e**; visual: *z*_57_=0.20, *p*=.845; motor: *z*_47_=0.20, *p*=.838).

Another measure that can provide an indirect measure of spectral quality is the residual error associated with fitting models to metabolite signals, i.e., fit error. If the fit error is high, this indicates that the model is poorly fit to the signal and can indicate poor signal quality. Thus, we also included the fit error associated with GABA+ and Glx in our regression analysis. However, we found no significant change in the correlation between GABA+ and Glx as a result of controlling for either GABA+ (**Fig. 2f**; visual: *z*_57_=0.03, *p*=.975; motor: *z*_47_=0.00, *p*=1.0) or Glx fit error (**Fig. 2g**; visual: *z*_57_=0.09, *p*=.929; motor: *z*_47_=0.03, *p*=.974).

It is challenging to rule out the possibility that signal quality contributed to the lack of a correlation between GABA+ and Glx. In particular, if GABA+ and Glx measurements were inaccurate due to poor signal quality, we would not expect them be correlated. However, if this were true, we would also expect them to be uncorrelated with other metabolite concentrations. To test this possibility, we computed the correlation between GABA+ and Glx and tCr and tNAA. We found that in visual and motor cortices both GABA+ and Glx were significantly positively correlated with tCr and tNAA (all *p*<.05), confirming that the measurements of GABA+ and Glx were sufficiently accurate to detect relationships between metabolites.

### Tissue composition

GABA is approximately twice as highly concentrated in grey matter compared to white matter (Petroff, Ogino, & Alger, 1988), and negligible in cerebrospinal fluid. Thus, inter-individual differences in the fractional tissue composition within the MRS voxel may have obscured a relationship between GABA+ and Glx. However, we found no significant change in the correlation between these neurotransmitters in visual or motor cortices as a result of controlling for the proportion of grey matter (**Fig. 2h**; visual: *z*_57_=0.03, *p*=.976; motor: *z*_46_=0.06, *p*=.953). Note that differences in voxel tissue composition may explain the difference in average GABA+ and Glx concentrations observed here and in Steel et al. (2020).

We next tested the possibility that differences in voxel tissue composition obscured a relationship between GABA+ and Glx by applying an alpha-tissue correction method (Harris, Puts, & Edden, 2015). However, we found that the correlation between these neurotransmitters in visual or motor cortices (visual: *r*=-.12, *p*=.364; motor: *r*=-.08, *p*=.573) did not change significantly as a result of applying this correction (visual: *z*_57_=0.16, *p*=.876; motor: *z*_46_=0.18, *p*=.854). Finally, we applied a tissue-correction method that accounts for differences in relaxation times across the tissue types within a voxel (Gasparovic et al., 2006). In line with the previous results, we found that the correlation between GABA+ and Glx in visual or motor cortices (visual: *r*=-.11, *p*=.425; motor: *r*=-.15, *p*=.322) did not change significantly as a result of applying this correction (visual: *z*_57_=0.08, *p*=.937; motor: *z*_46_=0.49, *p*=.628).

### Other neurochemicals

Previous work reported that the relationship between GABA+ and Glx in medial parietal cortex is moderated by the concentration of tCr and tNAA (Steel et al., 2020). While we found no significant change in the relationship between GABA+ and Glx as a result of controlling for tCr (**Fig. 2i**; visual: *z*_57_=1.16, *p*=.247; motor: *z*_47_=1.33, *p*=.182) or tNAA (**Fig. 2j**; visual: *z*_57_=0.69, *p*=.487; motor: *z*_47_=1.24, *p*=.214), we found a significant negative correlation between GABA+ and Glx in visual cortex, after controlling for the concentration of tNAA (*r*_58_=-.30, *p*=.021), and in motor cortex, after controlling for the concentration of tCr (*r*_47_=-.30, *p*=.040) and tNAA (*r*_47_=- .32, *p*=.027).

Steel et al. (2020) found no difference in the relationship between GABA+ and Glx in medial parietal cortex as a result of controlling for the concentration of Cho. We also found no significant change in the relationship between these neurotransmitters when we controlled for Cho (**Fig. 2k**; visual: *z*_57_=1.09, *p*=.274; motor: *z*_47_=1.24, *p*=.215); however, controlling for Cho, like controlling for tNAA resulted in a significant negative correlation between GABA+ and Glx in visual (*r*_58_=-.29, *p*=.026) and motor cortices (*r*_47_=-.30, *p*=.040). By contrast, although we found no difference in the relationship between GABA+ and Glx in visual or motor cortices as a result of controlling for the concentration of water (**Fig. 2l**; visual: *z*_57_=0.08, *p*=.934; motor: *z*_47_=0.57, *p*=.572), controlling for water did not result in a significant negative correlation in either area.

### Post-hoc Bayesian analysis

In contrast to previous work examining medial parietal cortex (Steel et al., 2020), we found no evidence for a positive correlation between GABA+ and Glx in visual and motor cortices. In a post-hoc analysis, we assessed the strength of the evidence for the null hypothesis (H_0_), where the alternative hypothesis (H_1_) is that there is a positive relationship between GABA+ and Glx. The results of the analysis are shown in **Table 2**. For the uncorrected GABA+ and Glx values, we find moderate evidence in favour of H_0_ for both visual and motor cortices (Jeffreys, 1961). For all corrected values, we find either moderate or strong evidence in favour of H_0_.

**Table 2.**
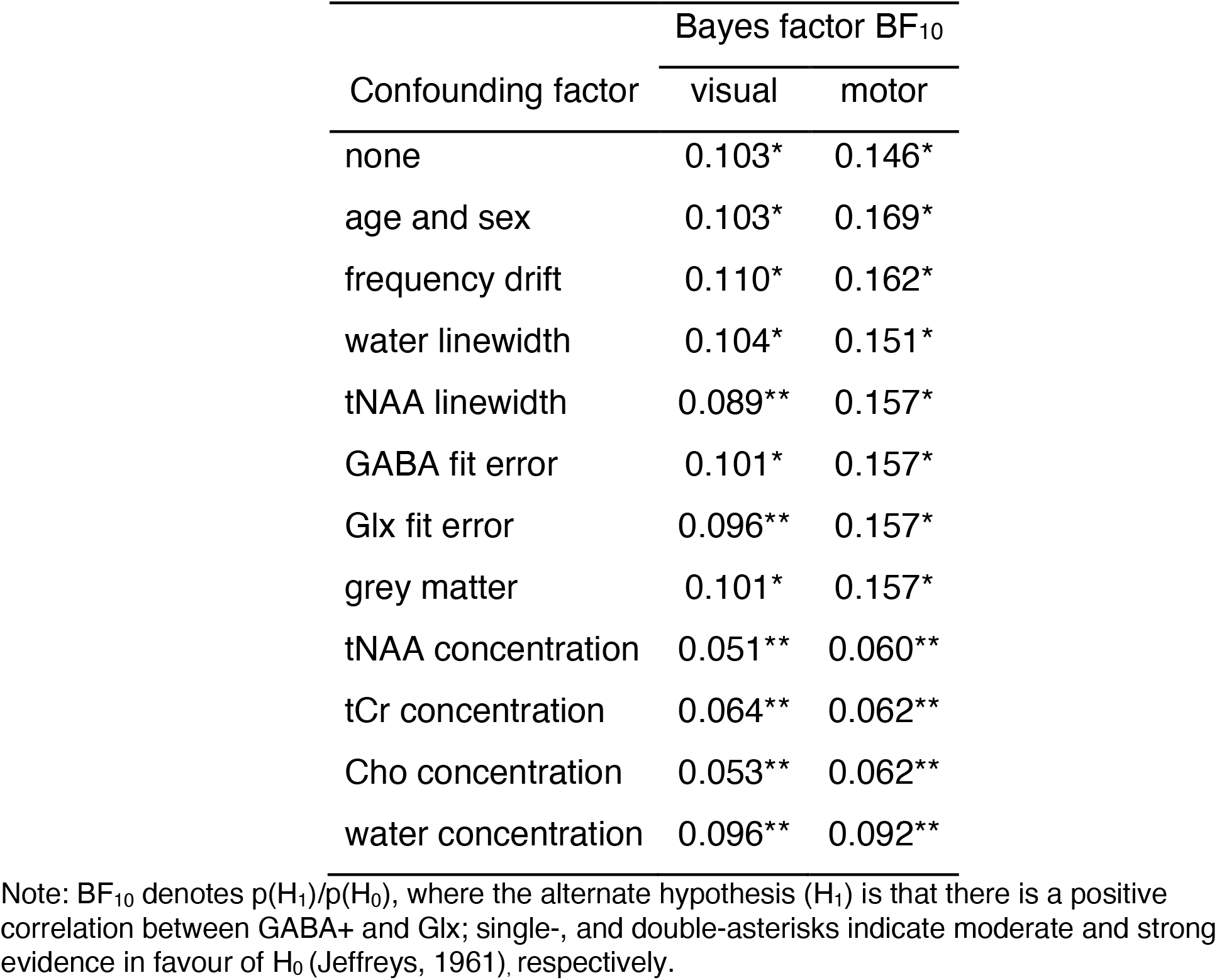
Probability of positive correlation between GABA+ and Glx

| Confounding factor | Bayes factor $BF_{10}$ | |
| --- | --- | --- |
|  | visual | motor |
| none | 0.103* | 0.146* |
| age and sex | 0.103* | 0.169* |
| frequency drift | 0.110* | 0.162* |
| water linewidth | 0.104* | 0.151* |
| tNAA linewidth | 0.089** | 0.157* |
| GABA fit error | 0.101* | 0.157* |
| Glx fit error | 0.096** | 0.157* |
| grey matter | 0.101* | 0.157* |
| tNAA concentration | 0.051** | 0.060** |
| tCr concentration | 0.064** | 0.062** |
| Cho concentration | 0.053** | 0.062** |
| water concentration | 0.096** | 0.092** |

Note: $BF_{10}$ denotes $p(H_1)/p(H_0)$ , where the alternate hypothesis ( $H_1$ ) is that there is a positive correlation between GABA+ and Glx; single-, and double-asterisks indicate moderate and strong evidence in favour of $H_0$ (Jeffreys, 1961), respectively.

### Visual stimulation control analysis

We found evidence that there is no positive correlation between GABA+ and Glx in visual and motor cortices. Consistent with previous work (Steel et al., 2020), spectra were acquired while participants were at rest. However, it is possible that a positive correlation between GABA+ and Glx in visual and motor cortices can only be observed during periods of relatively high activity. To test this possibility, we assessed the relationship between GABA+ and Glx in visual cortex while participants received visual stimulation (see the ‘mixed polarity’ condition of (Rideaux et al., 2019) for a detailed description of the visual stimulus). Consistent with the results from participants at rest, we found no evidence of a relationship between GABA+ and Glx in visual cortex of participants during visual stimulation (*r*_31_=.23, *p*=.213; **Fig. 3**).

**Figure 3.**
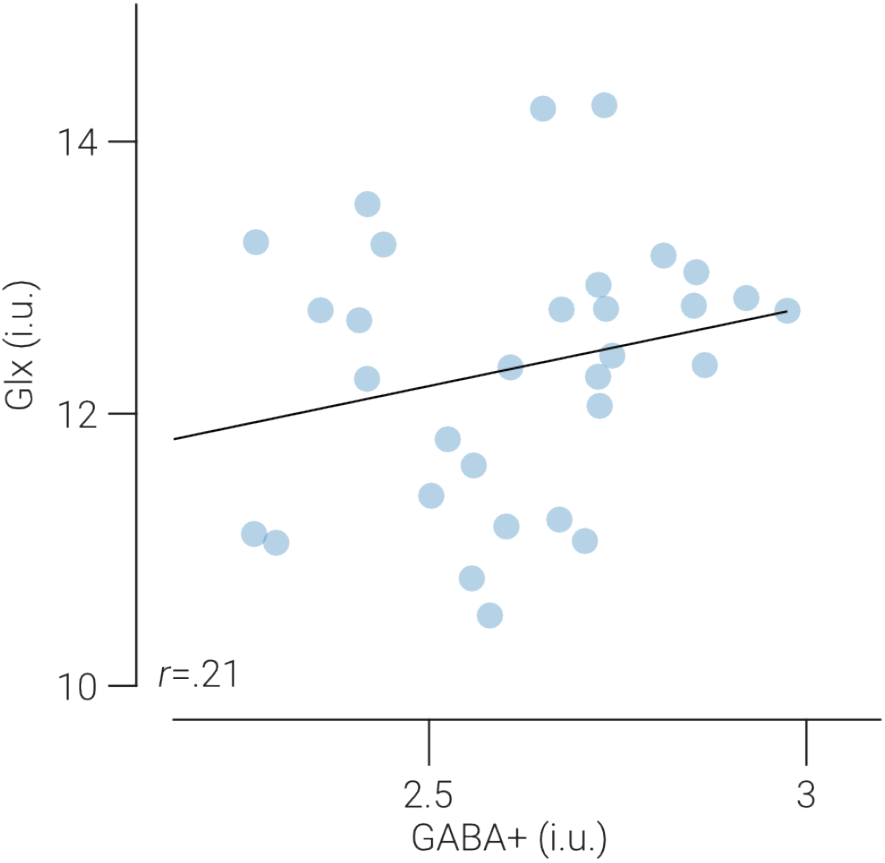
No balance between GABA+ and Glx in stimulated visual cortex. Glx concentration as a function of GABA+ concentration in visual cortices of participants while receiving visual stimulation. Glx and GABA+ values are referenced to water and expressed in institutional units (i.u.). Line indicates best linear fit.

## DISCUSSION

Previous MRS work revealed a positive correlation between GABA+ and Glx in medial parietal cortex of resting human participants (Steel et al., 2020). Here we performed the same test in visual and motor cortices and found no correlation between these neurotransmitters in either region. Post-hoc Bayesian analyses yielded moderate to strong evidence in favour of the null hypothesis, that is, there is no positive correlation between GABA+ and Glx. These results show that the relationship between these neurotransmitters is regionally localized, rather than a brain-wide characteristic. Consistent with (Steel et al., 2020), we controlled for age, gender, signal quality and tissue composition, and found no change in the relationship between GABA+ and Glx. However, controlling for the concentration of other neurochemicals (tCr, tNAA, Cho) revealed a negative correlation between these neurotransmitters in visual and motor cortices, suggesting a population-level imbalance in these regions.

These results appear to conflict with theoretical (Marín, 2012; Shadlen & Newsome, 1994; Van Vreeswijk & Sompolinsky, 1996; Vogels & Abbott, 2009) and empirical (Haider, Duque, Hasenstaub, & McCormick, 2006; Shu, Hasenstaub, & McCormick, 2003) evidence supporting canonical E/I balance within the brain. This discrepancy underscores differences in how E/I balance is measured. For example, in response to visual stimulation, electrophysiological work has shown concomitant excitatory and inhibitory activity (Anderson et al., 2000; Maffei & Turrigiano, 2008; Monier, Chavane, Baudot, Graham, & Frégnac, 2003; Priebe & Ferster, 2005; Xue et al., 2014), indicating balance, whereas MRS studies show GABA and Glu/Glx change in opposite directions (Kurcyus et al., 2018; Mekle et al., 2017; Rideaux, 2020; Rideaux et al., 2019), indicating imbalance. The negative correlation between GABA+ and Glx, and their changing in opposite directions, may be related to the GABA-glutamate cycle. In particular, at GABAergic synapses, Gln must be converted to Glu before being metabolised into GABA. Thus, increased GABA production could result in a temporary reduction in Gln and/or Glu, both of which comprise the Glx signal. These results do not undermine previous electrophysiological evidence for E/I balance. Rather, they demonstrate a dissociation between electrophysiologically measured I/E balance and the ratio of MRS visible GABA+ and Glx (the source(s) of which remain unclear).

Previous MRS work yielded inconsistent evidence for a balance between GABA+ and Glx in early visual and motor cortices. Some studies have reported a significant positive correlation between GABA and Glx/Glu in visual (Rafique & Steeves, 2020) or motor cortices (Rafique & Steeves, 2020; Terhune, Russo, Near, Stagg, & Cohen Kadosh, 2014), while others have failed to detect a relationship (Kurcyus et al., 2018; Terhune et al., 2014; Van Loon et al., 2013). These inconsistences may be a product of correlations based on relatively small sample sizes (Rafique & Steeves, 2020; Terhune et al., 2014) and less consideration of confounding factors (e.g., voxel tissue composition). Here we employed a larger sample size and directly tested whether or not there is a balance between GABA+ and Glx in visual and motor cortices by assessing the influence of confounding factors and applying Bayesian statistics that can provide evidence for the null hypothesis.

### Comparison of E/I observations in visual and motor cortices

The large body of evidence supporting E/I balance is primarily based on electrophysiological work measuring changes in activity occurring over time at relatively high spatial (e.g., single-cell recording) and temporal resolution (e.g., milliseconds) (Haider et al., 2006; Okun & Lampl, 2008; Shu et al., 2003); see (Isaacson & Scanziani, 2011) for a review. By contrast, we measured the average concentration of excitatory (Glx) and inhibitory (GABA+) neurotransmitters from a relatively large cortical region (e.g., 18 cm^3^) over a 13 min period and assessed their relationship across individuals. The dissociation between the previously established E/I balance in these areas and the concentration of MRS visible excitatory and inhibitory neurotransmitters may be explained by differences in spatial/temporal resolution or observations made within and between individuals. For instance, while there may be no population-level common ratio between GABA and Glu, there may be intraindividual balance that can be observed by measuring the concentration of these neurotransmitters over time. We recently assessed the relationship between change in GABA+ and Glx in visual cortex at low and high temporal resolution using a novel analysis that increased the temporal resolution of neurochemical estimates detected with MRS by a factor of five (Rideaux, 2020). We found a predictive relationship between these neurotransmitters that could only be observed at high temporal resolution. That is, we found that a change in GABA+ predicts an opposite change in Glx 2 min later. While this relationship does not suggest E/I balance, which would be evidenced by change in the same direction, the finding demonstrates that intraindividual relationships between these neurotransmitters can be detected with MRS, and further, that they can be obscured by the temporal resolution of the measurements.

### Differences between brain regions

The discrepancy between our findings and those of (Steel et al., 2020) may be explained by differences in information processing between medial parietal cortex and visual and motor cortices. The function of visual and motor cortices is relatively specialised. The region of visual cortex targeted by our voxel contains primary and intermediate visual cortex (i.e., areas V1, V2 and V3). These areas primarily support processing of visual input from the retina via the lateral geniculate nucleus (Hubel & Wiesel, 1959, 1968). Similarly, the cortical area targeted by our motor cortex voxel hosts the representation of hand and fingers and supports voluntary hand movement (Hluštík, Solodkin, Gullapalli, Noll, & Small, 2001; Penfield & Boldrey, 1937). Activity in sensorimotor areas may fluctuate more over time, due to natural variability in sensory inputs and motor activities (e.g., resting compared to running). In response to this variability, the E/I balance in these areas may be maintained at a higher spatial and/or temporal scale than that measured in the current study. For instance, there is an abundance of evidence for this balance when measured at higher temporal and spatial resolution, as measured with electrophysiological techniques (Anderson et al., 2000; Dehghani et al., 2016; Maffei & Turrigiano, 2008; Priebe & Ferster, 2005; Wilent & Contreras, 2005; Xue, Atallah, & Scanziani, 2014). By contrast, the medial parietal lobe acts as an information processing hub; it is associated with a diverse range of perceptual and cognitive functions, including (but not limited to) visual scene perception (Epstein, Parker, & Feiler, 2007; Silson, Steel, Kidder, Gilmore, & Baker, 2019), memory recall (Wagner, Shannon, Kahn, & Buckner, 2005), future prediction (Szpunar, Watson, & McDermott, 2007), and heading direction (Baumann & Mattingley, 2010). As such, its activity may be less variable and it may benefit from E/I balance maintained at a temporal and/or spatial scale that is detectable with MRS measurements averaged over a ∼10 min period.

Another, related, possible explanation is how these brain regions behave during rest. The medial parietal cortex is a core component of the default mode network (Andrews-Hanna, Reidler, Sepulcre, Poulin, & Buckner, 2010), and is active during rest and spontaneous thought (Fox, Spreng, Ellamil, Andrews-Hanna, & Christoff, 2015; Raichle, 2015). By contrast, visual and motor cortices are active during visual stimulation and motor movement, respectively, and relatively inactive while at rest. Thus, another possible explanation for the discrepancy in the relationship between GABA+ and Glx is that balance can only be observed during periods of relatively high neural activity. However, this explanation seems unlikely, as we found no evidence for a relationship between these neurotransmitters in visual cortex of participants who received visual stimulation. Further, electrophysiological work suggests excitatory and inhibitory activity in sensory cortices is balanced even during spontaneous activity (Atallah & Scanziani, 2009; Okun & Lampl, 2008).

### No clear link to metabolism

MRS-detected GABA and Glx/Glutamate is typically interpreted as indicator of inhibitory and excitatory neural activity (Edden, Muthukumaraswamy, Freeman, & Singh, 2009; Lunghi, Emir, Morrone, & Bridge, 2015; Stagg, Bachtiar, & Johansen-Berg, 2011a); however, the source(s) that give rise to the neurochemical signal that is detected remain unclear (Stagg, Bachtiar, & Johansen-Berg, 2011b). Steel et al. (2020) found that correlation between GABA+ and Glx was reduced when the concentration of tCr and tNAA, but not Cho, were controlled. The authors interpreted this as indicative of being related to metabolic function, as NAA is found exclusively within neurons (Urenjak, Williams, Gadian, & Noble, 1992) and Cr is involved in metabolic processes within multiple cell types (Andres et al., 2008); however, note that Cho is the precursor for the neurotransmitter acetylcholine (Nachmansohn & Machado, 1943). By contrast, we found that controlling for all three of these neurochemicals (tCr, tNAA, and Cho) had a similar effect on the relationship between GABA+ and Glx, producing more negative correlations (although controlling for tCr did not produce a significant correlation in visual cortex). Thus, our results suggest a different explanation, that is, that GABA+ and Glx share variance with all the neurochemical concentration measurements from the same spectrum. Here, and in Steel et al. (2020), the concentration of water was measured from separate (water unsuppressed) spectra and thus may have independent variance. Thus, while the results of Steel et al. (2020) appear are consistent with a link to metabolism, our results are more parsimoniously explained by factors that influence the entire spectra (e.g., participant movement).

### Clinical relevance

An E/I imbalance, in particular between Glu and GABA, is associated with a range of psychiatric and neurological illnesses including autism (Chao et al., 2010; Markram & Markram, 2010; Robertson, Ratai, & Kanwisher, 2016; Rubenstein & Merzenich, 2003; Vattikuti & Chow, 2010), schizophrenia (Kehrer et al., 2008), and epilepsy (Bradford, 1995; Olsen & Avoli, 1997). Evidence from MRS studies has revealed associations between Glx, GABA, and the severity of symptoms associated with autism spectrum disorder and schizophrenia (Brown et al., 2013; Egerton et al., 2012; Horder et al., 2013; Smesny et al., 2015), see (Foss-Feig et al., 2017) for a review. However, these associations appear to be regionally variable. For instance, autism spectrum disorder is associated with increased Glx in auditory cortex (Brown et al., 2013) and decreased Glx in basal ganglia (Horder et al., 2013). The discrepancy between the results from medial parietal cortex (Steel et al., 2020) and our findings from visual and motor cortices seem to be consistent with the regional variability reported in previous clinical MRS studies and highlight the diversity of relationships between neurochemicals, and neurochemicals and behaviour, across the brain (Greenhouse, Noah, Maddock, & Ivry, 2016).

### Limitations

Glutamate is the primary excitatory neurotransmitter in the central nervous system, whereas here we measured Glx, a complex comprising both glutamate and glutamine (Puts & Edden, 2012; Ramadan, Lin, & Stanwell, 2013). MRS work using phantoms indicates that glutamate contributes at least half the Glx signal in MEGA-PRESS difference spectra (Shungu et al., 2013; van Veenendaal et al., 2018), so it seems reasonable to interpret the concentration of Glx as representative of glutamate. However, it is possible that the contribution of glutamine to the Glx signal masked a correlation between GABA and glutamate. That is, a negative correlation between glutamine and GABA+ could have obscured a positive correlation between glutamate and GABA+, when combined in the Glx signal. Future work is needed test this possibility by directly measuring glutamate.

## Conclusion

We measured the concentration of GABA+ and Glx in visual and motor cortices and found moderate to strong evidence that there is no positive correlation between these neurotransmitters. These results contrast previous work that found a positive relationship between these neurotransmitters in medial parietal cortex and shows that there is no brain-wide interindividual balance between the concentration of GABA+ and Glx. Given that a balance was only found in one of three cortical locations, this suggests it may be the exception, rather than the rule. Our findings indicate a dissociation between the well-established E/I balance supported by previous theoretical (Marín, 2012; Shadlen & Newsome, 1994; Van Vreeswijk & Sompolinsky, 1996; Vogels & Abbott, 2009) and empirical evidence (Haider et al., 2006; Okun & Lampl, 2008; Shu et al., 2003) and the ratio between MRS-detected excitatory and inhibitory transmitters. This dissociation has implications for studies using this ratio as a proxy for E/I balance (Bang et al., 2018; Gu et al., 2019; Koizumi et al., 2018; Shibata et al., 2017; Takei et al., 2016), and for understanding and diagnosis of neurological and psychiatric illnesses characterised by an imbalance in excitation and inhibition.

## Acknowledgements

The work was supported by the Leverhulme Trust (ECF-2017-573), the Issac Newton Trust (17.08(o)).

## Notes

### Competing Interest Statement

The authors have declared no competing interest.

